# Conserving a urban habitat: Enhanced detection of common swift nests in buildings using a video fusion method

**DOI:** 10.1101/2025.08.25.671194

**Authors:** N. Sandoz, E. de Margerie

## Abstract

1. Common swifts (*Apus apus*) nest in cavities and cracks in building facades and roofs. It is a declining species in Europe, partly due to the loss of suitable breeding habitat caused by urban building renovation (e.g. thermal insulation). In several countries, building renovation projects imply quantifying the amount of habitat occupied by swifts that will be destructed, and proposing compensatory measures, such as installing nest boxes in the renovated building. A difficulty is that nesting swifts are uneasy to detect visually.
2. Here we report a new video method we used to improve swift nest detection. It is based on continuous filming of a building’s facade (for 1-2 hours), and later video-frame fusion to reveal swift flying in and out of cavities in the facade.
3. We succesfully used this method on two large building renovation projects, revealing larger than expected breeding populations (78 nests), and resulting in the implementation of ambitious compensation measures (212 nestboxes), commensurate with the habitat actually destroyed.
4. Based on a continuous dataset of swift passages during a full breeding season, we discuss favorable period in the season and day hours to efficiently detect nesting activity and assess building occupancy.
5. *Solution*. A video method is proposed to survey bird populations nesting in walls or cliffs. Video frame fusion reveals the fleeting movements of birds to their hidden nests. We provide software to use this tool in conservation projects, in particular ecological inventories prior to building renovation work.

## Introduction

The common swift (*Apus apus*) is a declining species in France (population decline estimated at -46% between 2001 and 2019; Fontaine et al., 2020) and in other European countries (e.g. -38% between 1995 and 2010 in the UK; Balmer et al., 2013). This species has a *Near Threatened* status in France (IUCN France, 2016). Among the possible causes of decline, studies have focused on climate change and climate anomalies, both in Europe (Thomson et al., 1996; Rajchard et al., 2006) and in the wintering area of the common swift (Ambrosini et al., 2011; Boano et al., 2020), located in sub-Saharan Africa (Akesson et al., 2020). Furthermore, as these birds are aerial insectivores, the decline in the quantity of insects available (Goulson et al., 2015; Hallmann et al., 2017), due in particular to the use of pesticides, is also considered to be a direct cause of the species’ decline (Moller et al., 2021). Lastly, as common swifts mainly nest in cavities and cracks in the facades and roofs of buildings, the gradual depletion of this urban habitat, due to the modernisation of buildings (e.g. renovation, waterproofing, external thermal insulation, demolition and new construction techniques) is a third cause of decline (Balmer et al., 2013; Genton & Jacquat, 2014; Crowe et al., 2010; Schaub et al., 2016; Dulisz et al., 2022).

When building demolition or renovation projects may impact this urban habitat, recommendations exist in several European countries for conserving or recreating habitat suitable for breeding swifts (e.g. Switzerland: Scholl, 2016; Ireland: Whelan et al., 2018; Italy: Gelati et al., 2019; France: LPO, 2024). In France, the destruction of common swift habitat is prohibited by law (as for any other protected species; Légifrance, 2025a,b), and can only be done after obtaining a derogation, which implies compensatory measures, in particular by installing nest boxes as part of the building project, and subsequent monitoring of nest box occupancy. Similar obligations exist in other countries, such as Germany and Poland (Schaub et al., 2016). The installed nest boxes can help keep common swift colonies in place, but it often takes several years for them to be occupied (Schaub et al., 2016; Dulisz et al., 2022).

The first step is to determine whether a building actually is a habitat occupied by swifts. An inventory of the number of cavities and cracks occupied by breeding birds on each façade of the building has to be carried out during one or more breeding seasons preceding renovation. This inventory is the basis on which the compensatory measures can be sized: in France, 2 to 3 nest boxes are usually required per active nesting cavity destroyed. The usual census method consists of visually observing the facade for at least 1 hour to detect swifts arriving or leaving their cavity (e.g. Schaub et al., 2016; Nobauer et al., 2018). In practice, this method requires a great deal of visual attention, as nesting swifts only pass through in a fraction of a second and are very discreet (contrary to non-breeding, screaming swifts). Even for experienced observers, this direct visual method is not infallible, particularly on large buildings where numerous cavities may be present. It is easy to confuse two neighbouring cavities if several passages occur at the same time, or if the observer’s attention is not full and constant.

## Methods

To overcome these difficulties, we developed a video method that increases the likelihood of detecting common swifts’ nests. First, we film a facade continuously for 1 to 2 hours from a fixed point. Second, to facilitate and accelerate the videos’ analysis back in the lab, we merge the frames of the video by applying a *minimum* filter, that retains only the darkest value of each pixel over the chosen number of frames (i.e. the chosen duration). This *video fusion* process sums up the duration of the video in a single still image, where the swifts’ passages appear clearly. This way, we obtain a record of all the birds passing through the wall during the recording session, with the precise locations of all the nesting cavities. (Fig. 1).

**Figure 1.**
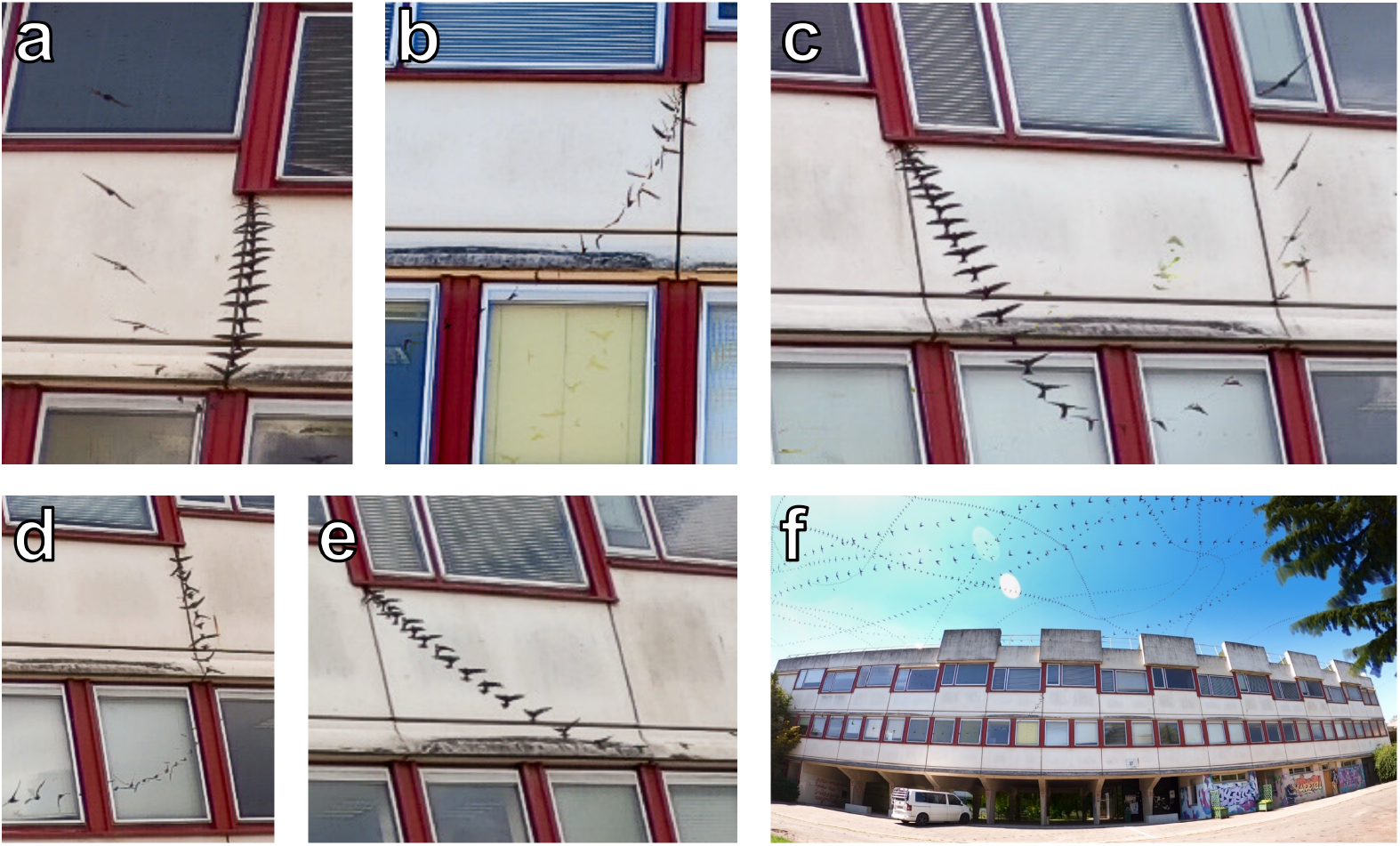
**(a-e)** Common swifts passing through cavities in the facade of a building **(f)**, filmed for 2 hours at 30 frames per second with a static wide angle camera (Gopro® Hero 8 black). Video frames were merged to produce a still image, by applying a filter that retains the darkest value of each pixel over the video duration. As swifts are darker than the building, their flight trajectories to and from cavities are revealed and easily detectable. Note that gliding flight (a, c, e) indicates a swift entering the facade, whereas flapping flight (b, d) indicates a bird exiting the building.

The fusion filter can be formally written as follows:

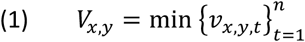

with:

- *v*_*x,y,t*_ the value of a pixel at position *x* (horizontally), *y* (vertically) and at time *t* in a video sequence (containing *n* frames).
- *V*_*x,y*_ the value of a pixel at position *x, y* in the final merged still image.

This filter is applied to each channel of the video file (i.e. 3 channels for RGB videos).

As storing all frames of a video in memory to find the overall minimum value is unpractical, the filter can be applied recursively while playing the video file:

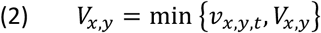

In practice, merging a very long video sequence into a single image is not the best option, because too many birds (and possibly insects) passing through the frame decreases the readability of the image, and can make it difficult to see whether the bird has really entered a cavity in the facade (i.e. the bird trajectory must disappear at the cavity entrance). Hence we usually merge the video by one-minute segments, which for a 2-hour video produces 120 still images, which are quick enough to inspect, and hence assess facade occupancy.

The favourable period to detect common swifts’ nests is when the adults perform regular roundtrips to feed the young (Arens, 2011a,b; Nobauer et al., 2018). In French Britanny, this usually corresponds to late June to mid July, but note that the period varies greatly depending on the latitude and year (Genton & Jacquat, 2014; Akesson et al., 2020). The usually recommended hours for visual detection are in the evening around sunset (Nobauer et al., 2018), but may also include the late morning (Schaub et al., 2016), where feeding visits are also frequent (Arens, 2011a,b). With the present video method, the efficiency of detection depends on the recording duration, as shown in Fig.2 : while a 1-hour video can be enough to detect a nest in the highest activity periods (i.e. late afternoons and evenings after hatching), a 2-hour video duration can provide an efficient detection throughout most of the day (10h-22h), for approximately 30 days after hatching.

**Figure 2.**
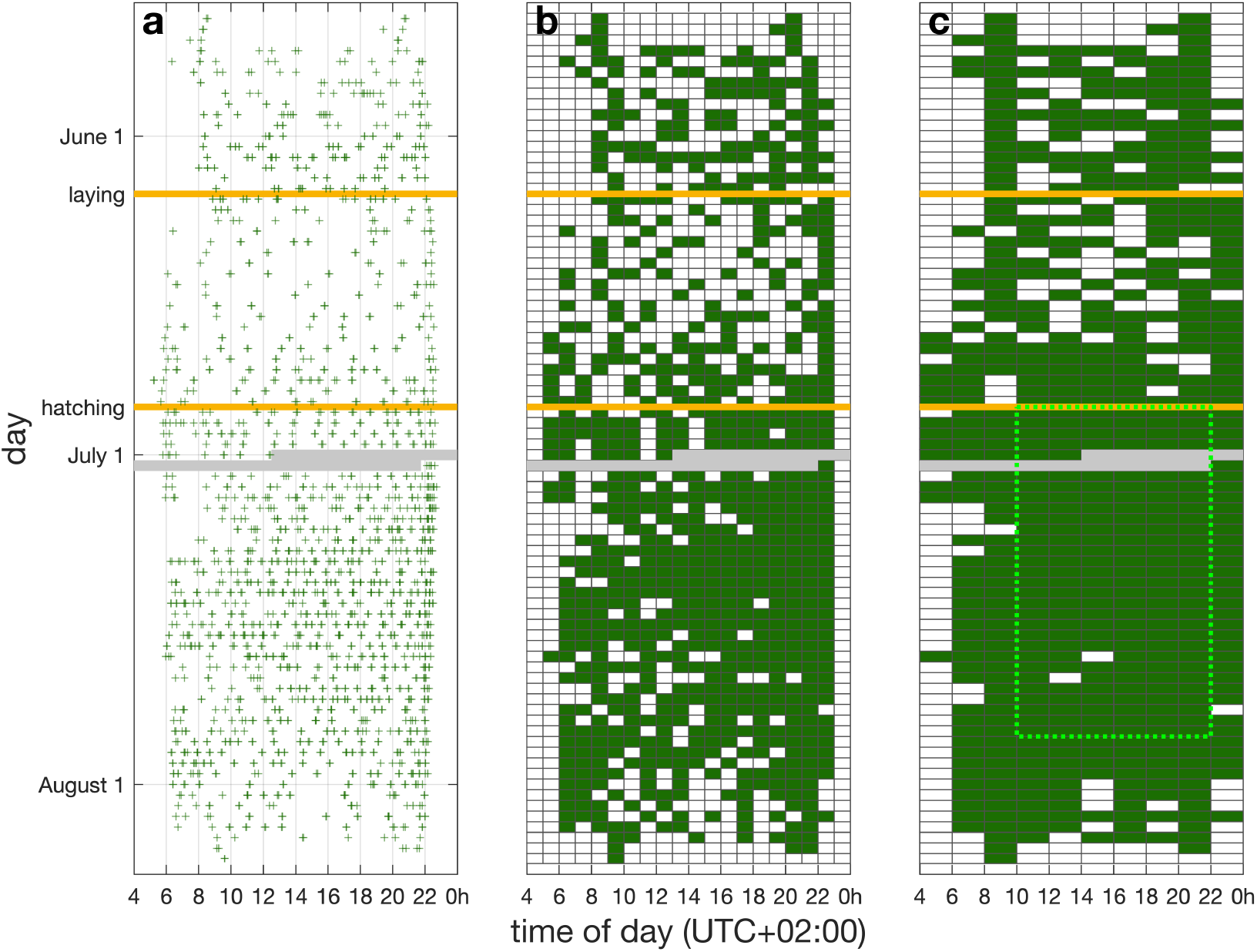
Example time distribution (days vertically downwards, hours horizontally) of passages performed by a couple of common swift adults flying in and out of their nest cavity. **(a)** Representation of each bird passage (green crosses) recorded using an IP camera placed inside a nest box in Rennes, France, from May to August 2024. Recording started on the first morning with both birds in the nest box (May 21). Two eggs were laid on June 7-8, and hatched on June 27-28. There was a network interruption on July 1-2, causing some data loss (grey area). Both adults and both juveniles left the nest box on August 7-8. Comparing the pre-laying period (May 21 – June 7), the incubation period (June 8 – 27) and the feeding period (June 27-August 8), the third period had the highest density of passages. **(b)** 1-hour time slots containing at least 1 passage are colored in green, while time slots containing no passage are left blank. Assuming an observer filming from outside the building for one hour continuously, the nest detection would have been most effective during the late afternoons and evenings (18-22h), as this period contains green slots almost enterily for approximately 20 days following hatching. **(c)** Following the same procedure with 2-hour time slots shows that doubling the video duration greatly improves passage recording probability: almost any 2-hour slot between 10h and 22h, for at least 30 days after hatching (green dotted perimeter) would have detected 1 passage (or more) in or out of the nest box.

## Results and Discussion

We applied this video method to survey common swift nests before two university campus renovation projects (external thermal insulation), in Rennes, France. By filming each facade of these large campus buildings for 2 hours, we were able to detect 78 active swift nests (de Margerie et al., 2023a,b), much larger populations than initially anticipated from occasional visual observations. Figure 1 displays the result of the *video fusion* filter for one of the campus building facade, and shows that split-second passages of swifts to their hidden nests are revealed and easily identified. As compensatory measures, the local biodiversity authority (DDTM 35) decided that 212 nest boxes should be integrated into the renovated facades, from 2024 to 2027.

Another benefit of the present method is that recorded images provide concrete evidence that birds inhabit a facade. Still images can easily be shared with the various parties involved in the building project, who may not have been able to see swifts entering the building for themselves.

To allow this video detection method to be used on other conservation projects, we provide a free software for merging video frames (https://github.com/edemargerie/videoFusion), with a detailed user manual and technical guidelines on how to implement the method in the field.

Note that the present method can be used to detect other bird species nesting in cavities in walls, roofs or cliffs (e.g. martins, sparrows, tits, redstarts), or even bats with some diurnal activity. For the frame fusion filter to yield easily interpretable images, the brightness value from the animal’s body needs to be contrastingly darker than the image background. In the case of a species appearing brighter than the background, one can modify filter (1) by replacing the *min* function with a *max* function.

## Author contributions

EdM conceived the ideas and designed methodology and software; NS and EdM collected the data; NS and EdM analysed the data; EdM led the writing of the manuscript. Both authors contributed critically to the drafts and gave final approval for publication.

## Acknowledgments

We would like to thank Xavi Bou, whose artistic photography work inspired the present method (https://xavibou.com/ornithographies/). We thank Hélène Demay and Justine Royer (LPO) for many discussions on common swift conservation in Rennes, France. We thank Paul Bernard (DMEAU) for accepting to perform early tests of our video technique and software. We thank Yuna Barbe, Léna Martin, Leslie Chauvin and Bertille Champion for their participation in video shooting and processing. We thank Ty Hedrick (North Carolina University, USA) for help on compiling the software. We thank Julien Le Bonheur for setting up and recording the nest camera footage. We thank Virginie Durier (CNRS) for proof-reading our manuscript.

## Data availability statement

*The swift nest data used to generate fig.2 will be available upon manuscript publication from the figshare data repository*.

## References

Åkesson S, Atkinson PW, Bermejo A, de la Puente J, Ferri M, Hewson CM, Holmgren J, Kaiser E, Kearsley L, Klaassen RHG, Kolunen H, Matsson G, Minelli F, Norevik G, Pietiäinen H, Singh NJ, Spina F, Viktora L, Hedenström A (2020) Evolution of chain migration in an aerial insectivorous bird, the common swift Apus apus. Evolution, 74, 2377–2391. 10.1111/evo.14093

Ambrosini R, Orioli V, Massimino D, Bani L (2011) Identification of putative wintering areas and ecological determinants of population dynamics of Common House-Martin (Delichon urbicum) and Common Swift (Apus apus) breeding in Northern Italy. Avian Conservation and Ecology, 6, 3. 10.5751/ACE-00439-060103

Arens H (2011a) Breeding biology of a pair of Swifts Apus apus ringed with an attached transponder. Vogelwelt, 132, 153–160.

Arens H (2011b) Transponder ringing for individual recording and observations of Swifts Apus apus. Vogelwelt, 132, 45–52.

Balmer D, Gillings S, Caffrey B, Swann B, Downie I, Fuller R (2013) Bird Atlas 2007-11: The Breeding and Wintering Birds of Britain and Ireland. BTO books. Thetford.

Boano G, Pellegrino I, Ferri M, Cucco M, Minelli F, Åkesson S (2020) Climate anomalies affect annual survival rates of swifts wintering in sub-Saharan Africa. Ecology and Evolution, 10, 7916–7928. 10.1002/ece3.6525

Crowe O, Coombes RH, Lysaght L, O’Brien C, Choudhury KR, Walsh AJ, Wilson JH, O’Halloran J (2010) Population trends of widespread breeding birds in the Republic of Ireland 1998–2008. Bird Study, 57, 267–280. 10.1080/00063651003615147

de Margerie E, Bernard P, Sandoz N (2023a) Détection vidéo des nids de Martinets noirs sur le campus Santé à Rennes (35). DMEAU. https://hal.science/hal-04302874v1

de Margerie E, Bernard P, Sandoz N, Chauvin L (2023b) Diagnostic écologique des bâtiments sur le campus de Beaulieu à Rennes (35). DMEAU. https://hal.science/hal-04302900v1

Dulisz B, Stawicka AM, Knozowski P, Diserens TA, Nowakowski JJ (2022) Effectiveness of using nest boxes as a form of bird protection after building modernization. Biodiversity and Conservation, 31, 277–294. 10.1007/s10531-021-02334-0

Fontaine B, Moussy C, Chiffard Carricaburu J, Dupuis J, Corolleur E, Schmaltz L, Lorrilière R, Loïs G, Gaudard C (2020) Suivi des oiseaux communs en France 1989-2019 : 30 ans de suivis participatifs. MNHN-Centre d’Ecologie et des Sciences de la Conservation, LPO BirdLife France-Service Connaissance, Ministère de la Transition écologique et solidaire. https://www.vigienature.fr/sites/vigienature/files/atoms/files/syntheseoiseauxcommuns2020_final.pdf

Gelati A, Ferraresi M, Gianella C, Ferri M, Poglayen G (2019) Progettare nel rispetto della protezione della BIODIVERSITÀ. Raccomandazioni e linee guida per la ristrutturazione e costruzione di edifici storici e moderni. Stazione Ornitologica Modenese, CISNIAR, Monumenti Vivi. https://www.asoim.org/doc/Piano%20emergenza%20e%20ricostruzione%20biodiversit%C3%A0%207-3-19%20versione%20stampa.pdf

Genton B, Jacquat MS (2014) Martinet noir: entre ciel et pierre. Cercle Ornithologique des Montagnes neuchâteloises, Musée d’histoire naturelle de la Chaux-de-Fonds. Editions de la Girafe.

Goulson D, Nicholls E, Botías C, Rotheray EL (2015) Bee declines driven by combined stress from parasites, pesticides, and lack of flowers. Science, 347, 1255957. 10.1126/science.1255957

Hallmann CA, Sorg M, Jongejans E, Siepel H, Hofland N, Schwan H, Stenmans W, Müller A, Sumser H, Hörren T, Goulson D, Kroon H de (2017) More than 75 percent decline over 27 years in total flying insect biomass in protected areas. PLOS ONE, 12, e0185809. 10.1371/journal.pone.0185809

Légifrance. (2025a). Conservation de sites d’intérêt géologique, d’habitats naturels, d’espèces animales ou végétales et de leurs habitats (Articles L411-1 à L411-3). Accessed 5 May 2025. https://www.legifrance.gouv.fr/codes/section_lc/LEGITEXT000006074220/LEGISCTA000006176521/#LEGISCTA000033035415

Légifrance. (2025b). Arrêté du 29 octobre 2009 fixant la liste des oiseaux protégés sur l’ensemble du territoire et les modalités de leur protection. Accessed 5 May 2025. https://www.legifrance.gouv.fr/loda/id/JORFTEXT000021384277/

LPO (2024) Rénovation du Bâti et Biodiversité. LPO. https://www.lpo.fr/la-lpo-en-actions/mobilisation-citoyenne/nature-en-ville/renovation-du-bati-et-biodiversite/renovation-du-bati-et-biodiversite-le-guide-technique

Møller AP, Czeszczewik D, Flensted-Jensen E, Erritzøe J, Krams I, Laursen K, Liang W, Walankiewicz W (2021) Abundance of insects and aerial insectivorous birds in relation to pesticide and fertilizer use. Avian Research, 12, 43. 10.1186/s40657-021-00278-1

Nöbauer S, Schmeller F, Griesberger P (2018) Evaluierung und Verbesserung einer Methodik zur Kartierung von Mauerseglern. Egretta, 56, 109–115.

Rajchard J, Procházka J, Kindlmann P (2006) Long-term decline in Common Swift Apus apus annual breeding success may be related to weather conditions. Ornis Fennica, 83, 66–72.

Schaub T, Meffert PJ, Kerth G (2016) Nest-boxes for Common Swifts Apus apus as compensatory measures in the context of building renovation: efficacy and predictors of occupancy. Bird Conservation International, 26, 164–176. 10.1017/S0959270914000525

Scholl I (2016) Sites de nidification pour les Martinets noirs et à ventre blanc. COMONE. https://www.ne.ch/autorites/DDTE/SFFN/faune/Documents/Travaux/Sites_Nidification_Martinets_Scholl_brochure_martinets_2016.pdf

Thomson DL, Douglas-Hhome H, Furness RW, Monaghan P (1996) Breeding success and survival in the common swift Apus apus: a long-term study on the effects of weather. Journal of Zoology, 239, 29–38. 10.1111/j.1469-7998.1996.tb05434.x

UICN France, MNHN, LPO, SEOF, ONCFS (2016) La Liste rouge des espèces menacées en France - Chapitre Oiseaux de France métropolitaine. Paris, France. https://inpn.mnhn.fr/docs/LR_FCE/UICN-LR-Oiseaux-diffusion.pdf

Whelan R, Hayes W, Caffrey B (2018) Saving swifts. BirdWatchIreland. https://birdwatchireland.ie/app/uploads/2019/10/Saving-Swifts-Guide_pdf.pdf

